# Assessing the blood-host plasticity and dispersal rate of the malaria vector *Anopheles coluzzii*

**DOI:** 10.1101/424051

**Authors:** James Orsborne, Luis Furuya-Kanamori, Claire L. Jeffries, Mojca Kristan, Abdul Rahim Mohammed, Yaw A. Afrane, Kathleen O’Reilly, Eduardo Massad, Chris Drakeley, Thomas Walker, Laith Yakob

**Author notes:** LY contact: Department of Disease Control, London School of Hygiene & Tropical Medicine, Keppel Street, London, WC1E 7HT.

## Abstract

Difficulties with observing the dispersal of insect vectors in the field have hampered understanding of several aspects of their behaviour linked to disease transmission. Here, a novel method based on detection of blood-meal sources is introduced to inform two critical and understudied mosquito behaviours: plasticity in the malaria vector’s blood-host choice and vector dispersal. Strategically located collections of *Anopheles coluzzii* from a malaria-endemic village of southern Ghana showed statistically significant variation in host species composition of mosquito blood-meals. Trialling a new sampling approach gave the first estimates for the remarkably local spatial scale across which host choice is plastic. Using quantitative PCR, the blood-meal digestion was then quantified for field-caught mosquitoes and calibrated according to timed blood digestion in colony mosquitoes. We demonstrate how this new ‘molecular Sella score’ approach can be used to estimate the dispersal rate of blood-feeding vectors caught in the field.

## Introduction

Although many disease vectors have demonstrable preference for a particular type of mammalian host to obtain a blood-meal, no insect vector of any of the major infectious diseases of humans or animals is exclusive to a single host species. Fifty years of host-choice studies have been conducted since the experiment in which Gillies released *Anopheles* mosquitoes into an enclosed space and compared the numbers flying into a room holding a human volunteer with those entering a room with a calf [1]. While useful, this type of experiment can only inform the intrinsic host preference of the vector which may or may not be indicative of what host species is bitten in natural field settings [2]. Many extrinsic as well as intrinsic factors play a part in who or what is ultimately bitten by a disease vector in a field setting and these have been summarised comprehensively [3]. Although it has been recognised for a long time that the same mosquito population will often adjust its biting towards a more locally available host species [4, 5], the extent to which this behaviour is plastic remains understudied even for the most important disease vectors [6].

Implicit to the spatial scale across which feeding choice changes is the vector’s dispersal ability. For example, if a vector tends not to disperse very far, a reasonable assumption may be that it will be less discerning in its choice of host and therefore be more likely to bite whatever is nearby. Of considerable hindrance to this field’s development is the absence of reliable methods for assessing disease vector dispersal ability. Conducting experimental studies on mosquito dispersal has been particularly challenging. The majority of such experiments has involved the mark-release-recapture of mosquitoes. However, the impact of handling mosquitoes combined with the typically low recapture rates – in the order of <2% for *An. gambiae* [7–11] – has limited what can be learned.

We investigate the blood-meal sources of *An. coluzzii* caught in traps situated within a range of alternative blood-host species availabilities in a malaria endemic village of southern Ghana. Uniquely, these data inform the spatial range across which this principle malaria vector adjusts its targeted blood-host species. Moreover, by matching these data with timed laboratory mosquito feeding experiments, we demonstrate how blood-meal digestion can be used to inform dispersal rates in a way that is broadly applicable to other haematophagous disease vectors. We discuss the potential that this experimental design has for studying the spread and control of vector-borne diseases.

## Results

To determine the spatial scale across which blood-host choice varies, a transect of six Centers for Disease Control and Prevention (CDC) resting traps with 50m spacing was set up outdoors. The transect extended from a cattle pen situated on the outskirts of Dogo village in southern Ghana to within an area of human residence. A total of 318 blood-fed *Anopheles* mosquitoes were collected over a five-night period. All but one of these were identified as *An. coluzzii* using a combination of species-specific PCRs and Sanger sequencing of a fragment of the ITS2 gene. The remaining insect was identified by ITS2 Sanger sequencing as *An. melas* and was excluded from the analysis.

The dominant mosquito blood-meal was of bovine origin with 73.8% of all meals being sourced from these hosts. Four (1.3%) individual mosquitoes were found to have solely fed on humans with an additional ten (3.2%) having a mixed feed of both bovine and human blood. Figure 1 shows how the bovine blood index (BBI) varied significantly across the transect, showing a decreasing trend with increasing distance from the cattle shed (OR=0.57 95%CI 0.47 – 0.71, p<0.01). The opposite trend was observed for human blood-meals with the HBI increasing significantly towards the village (OR=1.51 95% CI 1.05 – 2.17, p=0.027).

**Figure 1.**
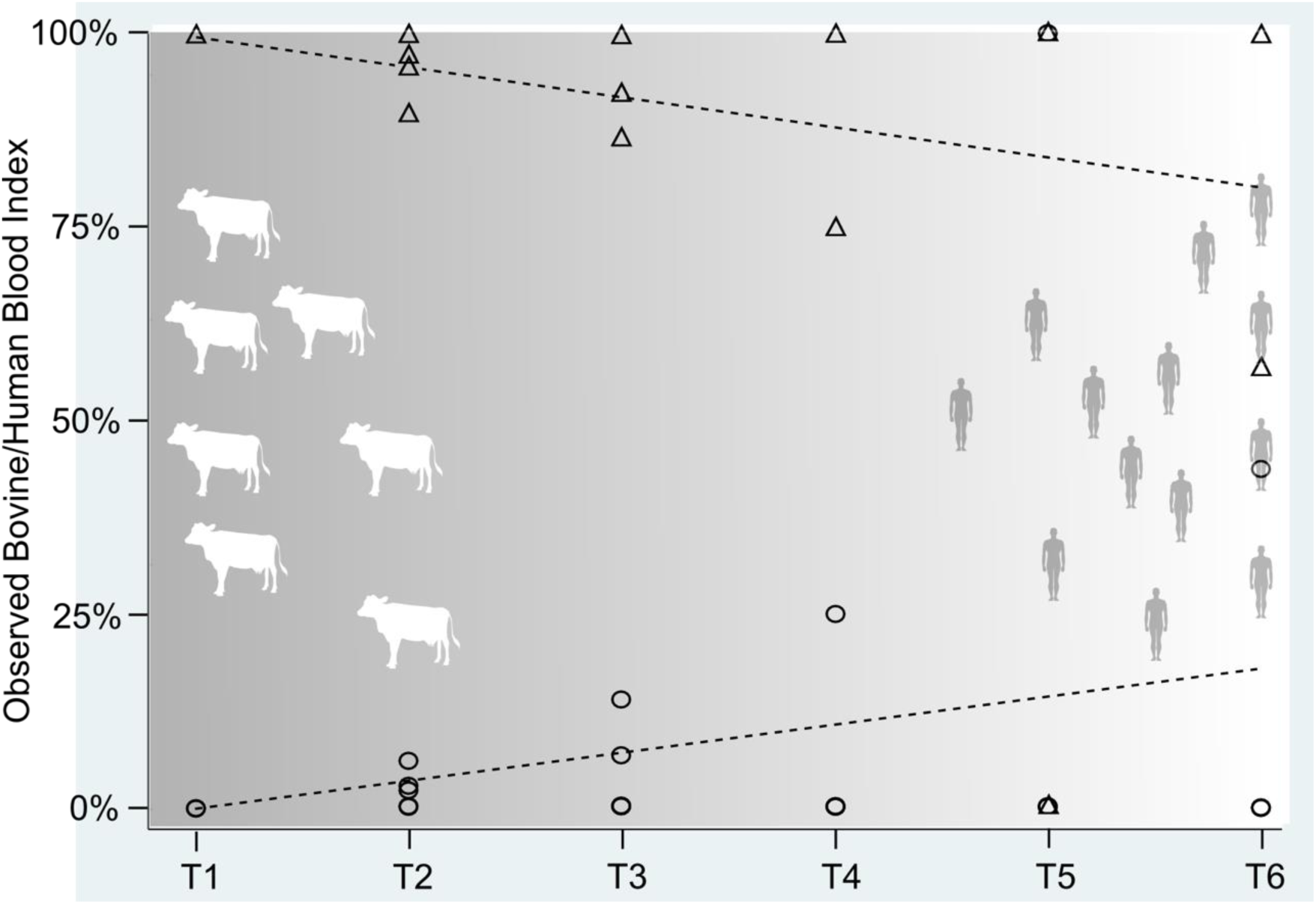
The human blood index (circles) and bovine blood index (triangles) for each nightly transect point (T1-T6) for all blood-fed *An. coluzzii* mosquitoes collected.

Measuring digestion of blood-meals in disease vectors has traditionally been accomplished through visual estimation – a method called Sella scoring [12]. Using quantitative PCR, we could eliminate the inherent subjectivity of this scoring while enhancing its resolution. Focusing on mosquitoes that had fed on cattle, it was observed that the quantity of blood-host DNA extracted from mosquitoes varied across the transect. Average PCR cycle threshold (Ct) values for bovine blood detection was 20.72 (95%CI 18.98-22.45) for mosquitoes caught by the cattle shed and 30.15 (23.14-37.16) for mosquitoes caught 250m away (p<0.01). To investigate this further, an experimental time series was performed with a laboratory colony of *An. coluzzii.* This allowed the effect of blood-meal digestion on Ct values to be investigated with the aim of producing mean Ct values for known time points post blood-feed. The time series showed Ct values increased with time post feed (p<0.01, see Figure 2). No bovine DNA was detected after the 60-hour time point. Serial dilutions of DNA extracted from bovine blood demonstrated the assay to have high levels of sensitivity and efficiency (E=-2.03, r^2^=0.97, slope= –3.26) with a detection limit equal to a 1000-fold dilution of DNA extracted from a freshly bovine blood-fed female *An. coluzzii* (Figure 2).

**Figure 2.**
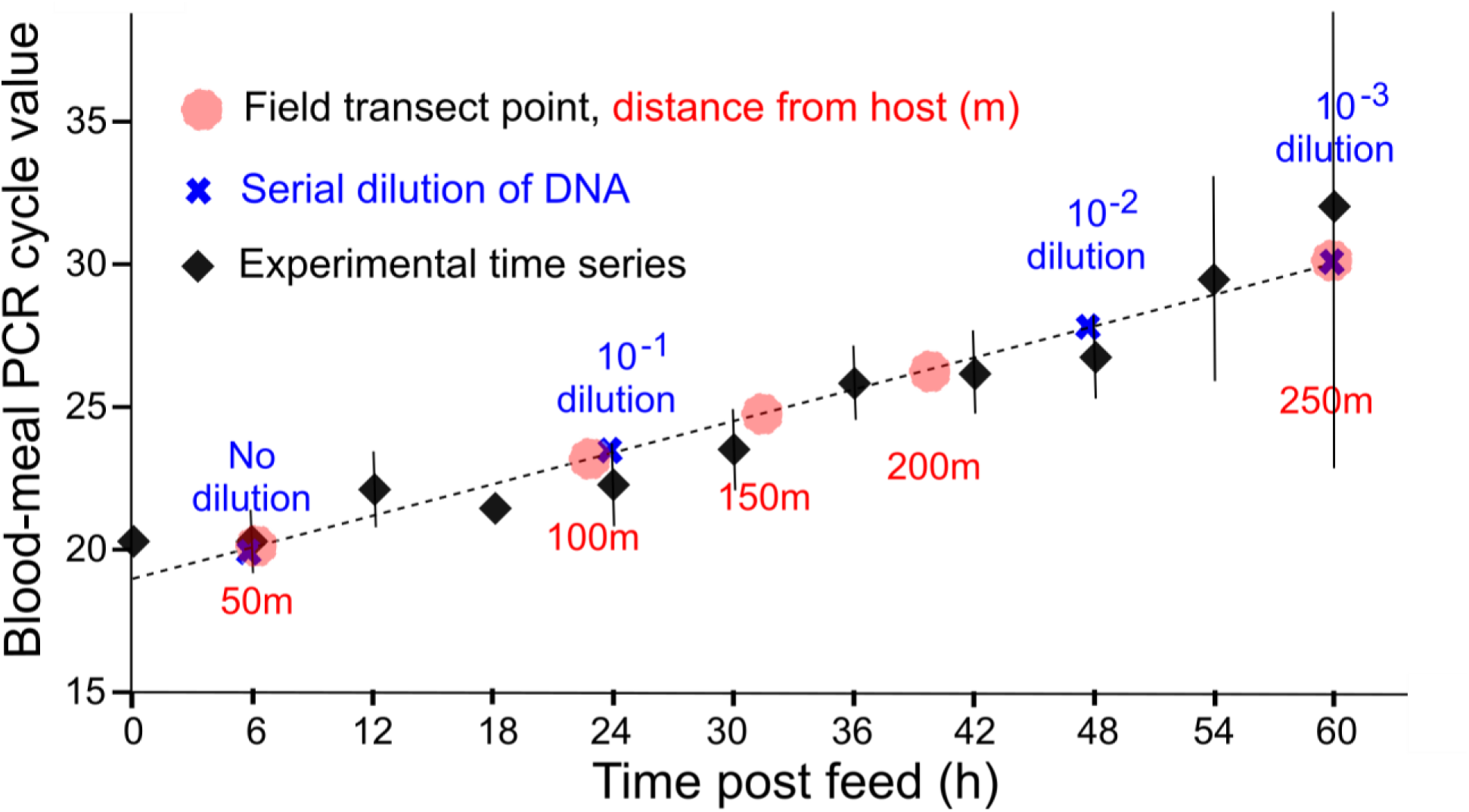
Effect of time post blood-meal on mean bovine Ct values produced from qPCR. Shown are the mean (bars indicate 95% CIs) of experimental time series (black), the serial dilution Ct values to assess assay sensitivity (blue), the mean Ct values of each transect point (red) and regression line used to predict time post feeding (dashed black line).

Regression analysis showed a positive correlation between bovine Ct value and time post feed in the experimental time series (r^2^ =0.92, slope = 0.183; see Figure 2). Calibrating the blood-meals of field-caught mosquitoes using the timed experiment with our mosquito colony, the dispersal rate of *An. coluzzii* could then be extrapolated: within 7 hours of feeding, mosquitoes typically remained within 50m of their blood-host but after 60 hours on average they had dispersed 250m (Figure 2).

As larger female mosquitoes typically obtain a larger blood-meal when feeding [13], we normalised for mosquito body size to exclude the possibility that the different quantity of bovine DNA across the transect was due to a corresponding trend in mosquito size. Ct values for bovine DNA were normalised against the Ct values for the corresponding mosquito ribosomal DNA (rDNA) gene used for species identification, producing a ratio of bovine (*Bos taurus* mtDNA)-to-vector DNA (An. *coluzzii* rDNA). A mean ratio was produced for each transect point and a significant positive correlation between the Ct ratio and distance from the cattle shed was retained (t=-2.1102, p<0.05).

## Discussion

Evidence for the strong influence of extrinsic factors on host selection was demonstrated through analysis of the blood-meals of *An. coluzzii* caught from the field using a novel sampling strategy. Using a transect of traps spanning a range of availabilities of different host species, significant differences were found for the host-choice of the same population of mosquitoes across a remarkably small spatial scale. This has significant implications for vector control. For example, initial field studies involving endectocidal applications on livestock have shown encouraging results in terms of long-lasting mosquitocidal effects [14, 15]. However, previously this strategy has only been considered for targeting zoophilic malaria vector species (e.g. *An. arabiensis*). Results presented here challenge this dogma; regardless of the local vector species, inclusion of endectocide-treated livestock could be justified and substantiated by replicating this relatively simple and inexpensive sampling method. Coupled entomological-epidemiological modelling frameworks already exist for using these data to inform projections of this novel vector control [16], including its use as part of an integrated vector management programme [17].

Linking the quantity of host-blood DNA isolated from mosquitoes caught at different distances from the host species with timed blood-meal digestion assays conducted on colonised mosquitoes presents a novel method for informing dispersal rates. Dispersal is recognised to underlie mosquito population structure [11] as well as human exposure to transmission [18] and our ability to control transmission [19]. Yet, knowledge of this critical aspect of behaviour has been hampered by our inability to produce reliable estimates of vector dispersal in the field. This study provides the first estimates using a non-intrusive method for measuring malaria vector dispersal that informs the mosquito’s dispersal rate across its feeding cycle (approximately 2.5 days). However, there are some limitations that require mentioning.

First, the numbers of mosquitoes captured nearby humans were lower than those caught adjacent to cattle; and while the numbers caught in 5 nights were sufficient to inform statistically significant trends across the transect, the variability between capture nights precluded our ability to infer the likely shape of dispersal (e.g., leptokurtic versus Gaussian). Future collections over longer periods should go some way to rectifying this and providing improved insight into the dispersal shape. Second, in order to estimate distances from blood-hosts these hosts must remain spatially confined. While this was possible in the current study because cattle were confined to their holding pen, the experimental design for conducting an equivalent experiment on human-blood digestion would need careful consideration. Third, blood-meal digestion levels of field-caught mosquitoes were calibrated with colonised mosquitoes maintained at a constant temperature and humidity. Realistic temperature/humidity regimens that better emulate natural diurnal patterns have sometimes been shown to significantly impact various aspects of mosquito metabolism [20]. Therefore, future experiments are required to ascertain the influence that fluctuating temperatures may have on blood-meal digestion.

Results presented in this study provide new insight into fundamental aspects of malaria vectors with important implications for malaria control strategy. Additionally, the novel experimental design presented offers a new paradigm in measuring dispersal that should be broadly applicable to other field-caught blood-feeding disease vectors.

## Methods

### Study site and mosquito collection

Mosquitoes were collected from the village of Dogo, in the Greater Accra region of Ghana (05°52.418 N, 00°33.607 E). The village is in the south-eastern coast of Ghana, with the Gulf of Guinea to the south and the Volta River to the east. The average rainfall is approximately 927 mm per year with main rainy season from April to June and a shorter second season in October. Temperatures range from 23 to 33 °C. The area is costal savannah with sandy soil, short savannah grass with some small/medium sized trees. The land is used extensively for grazing livestock as well as growing crops for local trade. Housing mostly consisted of concrete structures with concrete/brick walls and flooring. Some traditional mud style houses were also present, more so on the peripheral of the village.

Mosquitoes were collected across five consecutive nights in June 2017. The trapping set up consisted of CDC resting traps placed outdoors at 50m intervals forming a 250m transect comprising of six trapping points (denoted: T1 – T6). This transect was set beginning at an area of low human population density (T1, a cattle resting and overnight holding pen) and extending towards a human population (T6, the village of Dogo). Mosquitoes were collected overnight from 6pm to 6am.

Blood-fed mosquitoes were processed individually with transect location, night collected, genus and blood feeding status (determined using the Sella score) being recorded. Abdomens of blood-fed mosquitoes were pressed onto FTA^®^Classic cards (Whatman, GE Healthcare) to preserve the blood-meal for molecular analysis and the head and thorax were placed individually into wells of a 96 well plate. Excess blood fed mosquitoes were preserved in RNA later (Thermo Fisher Scientific Life Technologies) in a 96 well plate where necessary.

### DNA extraction

Mosquito abdomens were extracted individually. Samples were homogenised using a Qiagen TissueLyser II (Qiagen, UK) with a 5mm stainless steel bead (Qiagen, UK) placed in each sample tube in a 96 well plate format. Once homogenised, DNA was then extracted using the Qiagen DNeasy 96 kits (Qiagen, UK) following manufacturer’s protocol. Blood-meals preserved on FTA cards were punched out using a sterile steel 4mm radius punch (Bracket, UK). Resulting punches were incubated in ATL buffer and Proteinase K for 6 hours before DNA extraction was performed following manufacturer’s protocol. Extracted DNA was stored at –20°C until analysed.

### Mosquito species identification

Mosquito species identification was initiated using a real-time multiplex PCR assay targeting the rRNA gene [21]. Standard forward and reverse primers were used in conjunction with two species-specific Taqman probes. The reaction conditions were as follows: a 12.5μl reaction containing 1μl of genomic DNA. 6.25μl of Quantinova (Qiagen, UK) probe master mix. 800nM of forward and reverse primers (Thermo Fischer Scientific, UK), 200nM of *Anopheles arabiensis* probe (Sigma-Aldrich, UK) and 80nM of *Anopheles gambiae* probe (Applied Biosystems, UK). Samples were run on a Stratagene MX3005P (Agilent Technologies, USA) using cycling conditions of 10min at 95°C, followed by 40 cycles of 95°C for 25s and 66°C for 60s. The increases in fluorescence were monitored in real time by acquiring at the end of each cycle.

To differentiate between *Anopheles coluzzii* and *Anopheles gambiae* s.s. within the *An. gambiae* species complex a single end-point PCR was performed. This PCR targets the SINE200 retrotransposon and utilising an insertion in this area allows the two species to be distinguished following gel visualisation [22]. *An. coluzzii* produces a band at 479 bp with *An. gambiae* s.s. producing a band at 249 base pairs. Reaction was as follows: a 25μl reaction containing 0.5mM of forward and reverse primers (Forward:5’-TCGCCTTAGACCTTGCGTTA-3, Reverse:5’-CGCTTCAAGAATTCGAGATAC-3’), 12.5μl of Hot start Taq polymerase (New England Biolabs NEB, UK), 9.5μl of nuclease free water and 2μl of template DNA. Cycling conditions were as follows: 10min at 94°C followed by 35 cycles of 94°C for 30s, 54°C for 30s, 72°C for 60s,a final elongation step of 72°C for 10 minutes finished the cycling program.

PCR products were visualised on a 2% agarose gel using an Egel E-Gel iBase Power System and E-Gel Safe Imager Real-Time Transilluminator (Invitrogen, UK). The assay was performed on 10% of all samples identified as *An. gambiae* from the first assay with corresponding controls. Samples producing unknown or inconclusive results were sequenced (ITS2 Sanger sequencing) using primers originally developed by Beebe & Saul [23] and sequences were used to perform nucleotide BLAST (NCBI) database queries. PCR reactions were performed on a T100 Thermal Cycler (Bio-Rad Laboratories, UK) and amplified gene fragments were visualized by electrophoresis on a 2% agarose gel using an E-gel E-Gel iBase Power System and E-Gel Safe Imager Real-Time Transilluminator (Invitrogen, UK).

### Blood-meal identification

Samples were initially screened using bovine and human specific primers developed by Gunathilaka *et al* [24]. These primers where selected based on the abundance of available host species in the area. The reaction conditions consisted of a 10μl reaction including 0.5μM of forward and reverse primers (Integrated DNA Technologies), 5μl of SYBR green master mix (Roche, UK), 2μl of nuclease-free water (Roche, UK) and 2μl of template DNA. PCR was run on a LightCycler 96 real-time PCR machine (Roche, UK) under the following cycling conditions: pre-incubation of 95°C for 600s, 40 cycles of 95°C for 10s, 62°C for 10s and 72°C for 30s followed by a melting analysis.

Human positive blood-meals (including potential mixed feeds) from the above assay were confirmed using the Promega Plexor^®^ HY Human DNA forensic detection kit (Promega, UK). Assay was performed following manufacturer’s protocol using a Stratagene MX3005P (Agilent Technologies, USA) real-time PCR machine.

### Lab assessment of blood-meal DNA degradation rate

Approximately 500 female *An. coluzzii* mosquitoes (N’gousso strain originally collected from Yaounde, Cameroon) were placed into an insect cage (Bugdorm, Watkins and Doncaster, UK) and, using a Hemotek, fed for 15 minutes on bovine blood collected from a UK based abattoir (First Line UK (Ltd), UK). Mosquitoes were reared at the London School of Hygiene & Tropical Medicine under standardized conditions in an incubator (27°C and 70% humidity with a 12:12 light/dark cycle) and given access to 10% sugar solution. Female mosquitoes were individually collected and checked for feeding status. Only overtly fully fed mosquitoes were selected for the experiment. Fully fed females were separated into paper cups covered with netting; each cup contained a maximum of 30 female mosquitoes. Every 6 hours a single cup was removed and placed in a - 80°C freezer to kill the mosquitoes and stop blood-meal digestion. This was repeated until the mosquitoes had completely digested the blood-meal or were visually gravid. DNA was extracted using the above protocol from seven whole bodies for each time point. A 1:10 serial dilution of all time = 0 samples was used to generate standard curve with dilutions being made down to 1 × 10^-7^. The standard curve was used to assess assay sensitivity (limit of detection) with the resulting Ct values from each time point being used to estimate the time post blood-feed for the field-caught mosquitoes. Both species identification and blood-meal source were confirmed molecularly using the protocols stated above. Trends in blood indices across the transect were tested for the field-caught mosquitoes using logistic regression.

## Acknowledgements

JO has an MRC London Intercollegiate Doctoral Training Partnership Studentship. TW and CLJ are funded through a Wellcome Trust/Royal Society Sir Henry Dale Fellowship (101285/Z/13/Z) awarded to TW. LY received funds from a Royal Society Research Project (RSG\R1\180203).

## Competing interests

The authors declare that there are no competing interests

Statement of authorship: LY designed the study. JO, CLJ, MK, ARM, YAA, TW conducted the experiments. LFK, KOR, LY analysed the data. JO, LFK, CLJ, MK, ARM, YAA, KOR, EM, CD, TW, LY interpreted results and contributed towards drafting the manuscript.

